# Real-cyber hybrid neural network for predicting neural circuits in mouse decision-making

**DOI:** 10.1101/2024.10.15.618576

**Authors:** Yutaro Ueoka, Hayato Maeda, Shuo Wang, Akihiro Funamizu

**Affiliations:** Institute for Quantitative Biosciences, The University of Tokyo, Laboratory of Neural Computation, 1-1-1 Yayoi, Bunkyo-ku, Tokyo 113-0032, Japan; Department of Life Sciences, Graduate School of Arts and Sciences, The University of Tokyo, 3-8-2, Komaba, Meguro-ku, Tokyo 153-8902, JAPAN

## Abstract

A major challenge in neuroscience is to elucidate how the brain processes sensory input to generate behavior, especially given the difficulty to measure neural activity across the whole brain. To address this limitation, previous studies have used artificial neural networks (ANNs) and modeled decision-making circuits in the brain. Here, we developed a real-cyber hybrid neural network (HNN) that directly input the activity of real neurons to an ANN, imposing biological constraints on the model. Using spike inputs from cortical or subcortical regions, the HNN predicted body movements of head-fixed mice during a task better than the ANN did. This improvement arose not only from the added mouse neuronal activity, but from the activity generated within the HNN. The generated activity potentially included unrecorded neuronal activity from mice, which was difficult for the ANN. We propose that HNNs develop brain-like activity to predict animal movements and compensate for unrecorded neural activity.

## Introduction

A fundamental function of the brain is making decisions based on sensory inputs. These decisions often accompany the physical body movements of an organism on a continuous time scale^1^. To understand the neural circuit involved in this decision-making process, previous studies have used brain-wide neural recording with two-photon microscopy^2–6^ or silicon probes^7–10^ to identify the neural correlates of essential task features, such as sensory inputs, action selection, and internal variables of the brain, such as reward expectations^7,11^. However, there are limitations to recording neuronal activity in the whole brain simultaneously, making it difficult to understand how the brain drives body movements for decisions.

In parallel with physiological studies, artificial neural networks (ANNs) have become successful tools for modeling how the neural circuits of the entire brain drive decision making^1,12,13^. ANNs can perform different strategies^13,14^ and often model simplified actions to understand the computational theory of decisions^15^. In addition to focusing on discretized choices, recent studies have used ANNs to investigate how the brain generates physical body movements on a continuous time scale to make decisions^1,16^. These studies showed that the activity of artificial units and that of neurons are somewhat correlated^2,4,12,17^, suggesting that the network model provides a hypothesis of how the brain moves the body on a continuous time scale.

Can we develop a brain-like network model in which the artificial units of ANNs exhibit activity patterns similar to those of the real neurons of animals? Although ANNs have become important tools for modeling brain networks^18^, limitations have also been reported^19^. Here, we propose a real-cyber hybrid neural network (HNN) that integrates the artificial units of an ANN and the real neurons of mice (**Fig. 1a**). Our HNN is a recurrent neural network based on reservoir computing (RC), in which only the readout-layer weight is trained to predict mouse behavior from time series of task features and neural activity. We chose RC, which does not train the internal layers, to test the idea of the HNN in a simple and interpretable model focused on neural activity propagation. Neural activity, recorded from mice, was input into the RC and propagated to constrain the activity of artificial units. We used the dataset from our previous study in head-fixed mice^20^, in which we used a Neuropixels 1.0 probe to electrophysiologically record neural activity from the orbitofrontal cortex (OFC), motor cortex (M1), parietal cortex (PPC), auditory cortex (AC), striatum (STR) and hippocampus (HPC) during a tone-frequency discrimination task.

**Figure 1.**
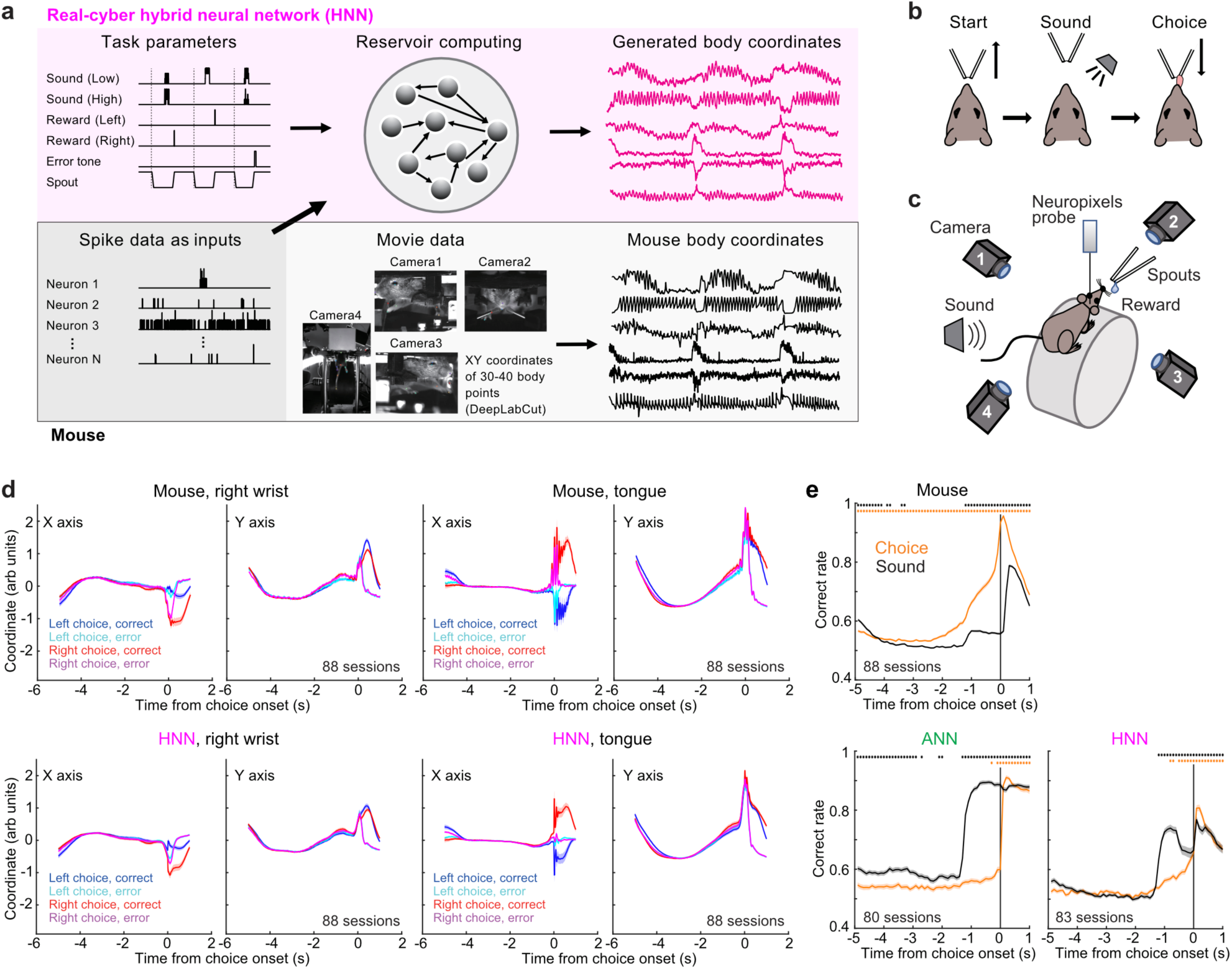
Real-cyber hybrid neural network (HNN) **a.** Schematic of the HNN. Six task inputs (sound (low), sound (high), reward (left), reward (right), error tone, and spout) and the spike data from each session were used in the HNN. **b.** Scheme of the tone-frequency discrimination task in head-fixed mice in our previous study^20^. The trial started by moving the spouts away from the mouse. After a sound presentation, the spouts were presented to the mice, and the mice licked the left or right spout to receive a reward. **c.** Task setting with 4 video cameras. **d.** XY trajectories of body points in mice and outputs of the HNN. The average trajectories of the right wrist and tongue are shown (means and standard errors, 88 sessions from 6 mice). **e.** Choice and sound category decoding from body trajectories. The choices and sounds were decoded every 100 ms. The Wilcoxon signed-rank test was used to investigate whether the decoding performance with cross validation was greater than 0.5 (means and 95% confidence intervals, *: p < 1e-10; 88, 80 and 83 sessions from 6 mice for the Mouse, ANN and HNN graphs, respectively).

We found that the HNN outperformed both conventional machine learning models and an ANN with only artificial units in predicting mouse body movements. This improvement was achieved by spike inputs from almost all the recorded brain regions, except the HPC. We found that the HNN, similar to real mouse, predicted choice-dependent body movements even before the mice chose the left or right spout. These early choice-dependent movements were not reproduced by the ANN. Further, when we developed an ANN and employed neural activity separately in parallel for prediction, we were unable to match the predictive accuracy of the HNN, indicating that inputting neural signals into the HNN contributed to generating artificial unit activity that enhanced prediction performance. When we trained HNN without using a part of recorded neurons, the artificial units in HNN had similar activity to the removed neurons compared to the ANN. Similarly, when we trained the HNN without sound-responsive neurons, the artificial units were still active at the sound onset. These results suggest that the HNN develops brain-like activity to predict animal body movements by propagating spike inputs through a recurrent network and compensating for unrecorded neuronal signals.

## Results

### A real-cyber hybrid neural network (HNN) predicts mouse body movements using task inputs and spike activity

In this study, we developed an HNN that uses the activity of real neurons in mice, as well as the task parameters, as the inputs of a recurrent neural network using reservoir computing (RC) (**Fig. 1a**). To develop the HNN, we used the data from our previous study, in which neuronal activity was electrophysiologically recorded with Neuropixels 1.0 during a tone-frequency discrimination task (**Fig. 1b, c**)^20^. In the task, the mice were head fixed and placed on a cylinder treadmill. In each trial, a sequence of low- or high-frequency pure tones of tone cloud was presented^21–23^. The reward site was either a left or right spout depending on the dominant frequency of the tone cloud (**Methods**). We recorded neural activity from the OFC, M1, PPC, AC, STR, and HPC. The simultaneously recorded neural activity from each brain region was used to train the HNN.

During the task, the body movements of the mice were captured by 4 or 5 video cameras with sampling rates of 137 and 140 Hz in 53 and 35 sessions, respectively (**Fig. 1c**). We extracted the XY trajectories of 30 or 40 body points in the 4- or 5-camera settings with DeepLabCut (DLC)^24^. Using the data collected during the task, the HNN generated the predicted XY trajectories of these body points on a continuous time scale based on the inputs of task parameters (i.e., sounds, rewards, and spout movements) and simultaneously recorded spike data. We trained the HNN using the initial 80% of the data from each session. The RC units were randomly connected based on a set of given hyperparameters. The hyperparameters of RC (i.e., number of nodes, expected number of connections per node and leaking ratio) were selected to achieve the optimal performance with the training data (**Fig. S1**). We then tested the performance of the HNN with the remaining 20% of the data (**Methods**).

We first analyzed the XY trajectories of body points extracted with DLC (**Fig. 1d and S2**). To standardize the frames per second (fps) across all the sessions, the sessions with 140 Hz were downsampled to 137 Hz via linear interpolation. We found that the body trajectories of the mice depended on the sound category (low or high), choice (left or right), and outcome (reward or error). The body movements generated by HNN varied depending on the task parameters. Using the body movements of the mice and the movements generated by the HNN, we decoded the choices and sound categories via sparse logistic regression with cross validation (**Fig. 1e**) (**Methods**). In mice, body movements predicted the left or right choice in each trial, and the performance of sound decoding was worse than that of choice decoding. The choice decoding performance in the HNN exceeded the chance level of 0.5 starting at 0.8 s before the choice. To evaluate the contribution of neural inputs, we developed an ANN that also used the RC architecture but excluded neural inputs. We also decoded the choice and sound categories using body trajectories generated by the ANN without spike inputs. The choice decoding accuracy of the ANN began to improve at 0.3 s before the choice. In contrast to the ANN, which receives only task parameters and has difficulty predicting the mice’s incorrect choices in advance, the HNN receives real neural activity as input, allowing it to generate choice-dependent body movements similar to those of the mice.

### The HNN outperforms both the ANN without spike inputs and other machine learning models in predicting mouse body movements

To validate the performance of RC architecture, we developed two models to estimate mouse body movements (**Fig. 2a**) (**Methods**). The average model (AM) used the mean trajectory of body coordinates in the training data. The generalized linear model (GLM) estimated the body coordinates with linear combinations of 420 task-parameter kernels (i.e., sound, reward/error, and spout movements). We analyzed the root mean squared error (RMSE) between the generated and actual body coordinates in 88 sessions from 6 mice. For all models, we separated the data into an initial 80% (training set) and a final 20% (test set). The RMSE was defined as the average RMSE of all the XY trajectories of body points (60 or 80 XY trajectories for 4 or 5 camera settings) in the test data. We found that the RMSEs of the ANN and HNN were smaller than those of the AM and GLM (**Fig. 2b**). These results suggest that RC architecture is suited for the behavior prediction.

**Figure 2.**
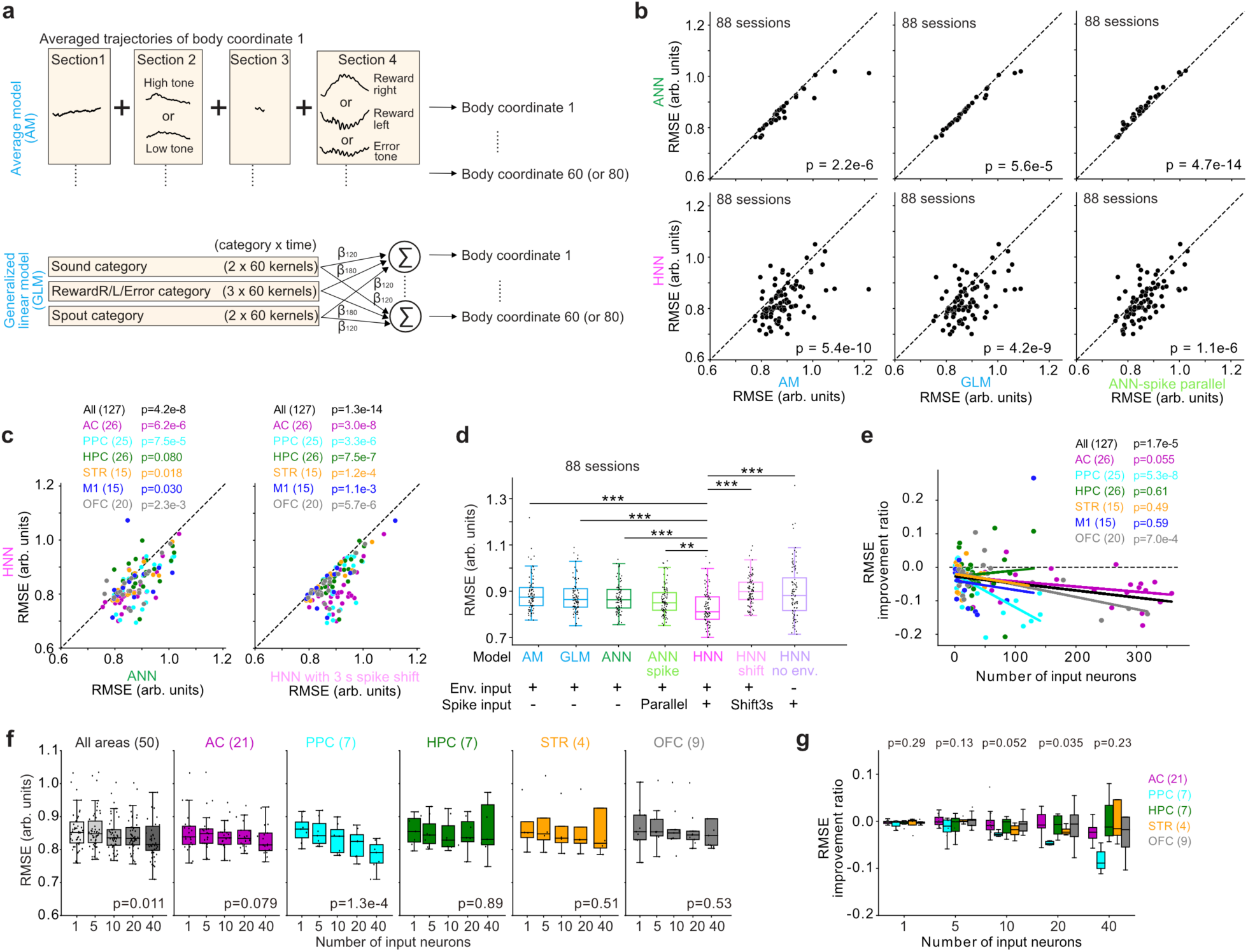
Performance of the real-cyber hybrid neural network (HNN) **a.** Scheme of the average model (AM) and generalized linear model (GLM). The AM calculates the mean trajectories of each body point in the training data (the first 80% of all the data) divided into four sections: (1) before tone onset, (2) between tone onset and spout arrival, (3) between spout arrival and choice, and (4) after choice. We compared the body trajectories in the test data (the last 20% of all the data) with the mean trajectories in the training data to compute the root mean squared error (RMSE) at each time point. The GLM used 60 kernels for each task category (**Methods**). Sound kernels started from tone onset (section 2) and lasted 6 s with steps of 0.1 s (60 kernels). The reward and error kernels started from the onset of the outcome and lasted 6 s (section 4). Spout kernels started at the arrival and departure of spouts and lasted 6 s (sections 1 and 3). We trained the GLM with the first 80% of all the data and fit it to the last 20% of all the data **b.** RMSEs of the HNN and ANN in each session compared with those of the AM, GLM and ANN-spike parallel (ANNSP) (Wilcoxon signed-rank test in 88 sessions from 6 mice). **c.** Improvements in performance in reservoir computing (RC) with precise spike inputs. Left: RMSEs of RC with and without spike inputs (HNN and ANN). Right: RMSEs of HNN using precise spike inputs and 3-s shifted spikes (HNN and HNN shift). The neural activity from the PPC-HPC or STR-M1 was simultaneously recorded with one Neuropixels probe in 24 or 15 sessions, respectively (39 sessions in total out of 88 sessions). The performance of the HNN using spike inputs from each brain region was independently analyzed (plots show the data from 88 + 39 = 127 sessions in total from 6 mice) (**Methods**) (Wilcoxon signed-rank test). **d.** Summary of HNN performance. HNN no env. used only the spike data as the inputs of RC (**: p < 1e-5, ***: p < 1e-7 in Wilcoxon signed-rank test, 88 sessions from 6 mice). The central line and edges in the boxplots represent the median, 25th, and 75th percentiles, respectively; the whiskers represent most extreme data points inside the limits of 1.5 times the interquartile range, here and throughout the paper. **e.** Relationship between the number of spike inputs and the performance of the HNN. The RMSE improvement ratio was the relative change in the RMSE of the HNN compared with that of the ANN (**Methods**). Robust linear regression was used to analyze the significance of the slope (t test). **f.** Relationship between the number of neurons and the performance of the HNN within sessions. Sessions with 40 neurons or more were used. A total of 1, 5, 10, 20, or 40 neurons were randomly chosen from each session. Since only 2 sessions had data from M1, we did not show the individual plots of M1. Robust linear regression was used to analyze the significance of the slope (p values from the t test). Outliers with RMSE improvement ratios > 0.15 were excluded from the figures. For **f** and **g**, plots including all the data points are shown in **Fig. S3**. Boxplots are the same as those in **d**. **g.** Comparisons of RMSE improvement ratios across brain regions for each number of neurons (p values from one-way ANOVA). Outliers with an RMSE > 1.1 were excluded from the figures.

We next validated the effect of spike input to HNN. We compared the performance of the HNN and ANN (**Fig. 2c, 2d**). The RMSEs of the HNN were smaller than those of the ANN, indicating that the spike inputs improved the estimation of body movements. We further found that the performance of the HNN was better than that of the HNN with 3-second-shifted spike inputs (**Methods**), suggesting that the improvement in performance was not achieved simply by the presence of spike-like inputs or their spectral properties. Rather, it suggests that the precise timing of spikes is crucial for generating body movements. We also found that the prediction accuracy of the HNN was better than that of the HNN without task-parameter inputs, indicating that neural activity inputs alone were not sufficient to generate accurate body movements (**Fig. 2c, 2d**). These results suggest that the inputs of both neural activity and task parameters are essential for generating mouse body movements over a continuous time scale.

To investigate whether the improvement in prediction accuracy comes from the neural activity itself or from processing the spike inputs as unit activity within the RC network, we constructed a standard ANN and placed the spike inputs in parallel with the ANN units—without providing them to the RC—to create the ANN-spike-parallel (ANNSP) model. The predictive accuracy of ANNSP was higher than that of the ANN, but lower than that of the HNN (**Fig. 2b**). This result suggests that the internal processing of spike inputs in the RC contributes an additional boost in prediction performance.

Next, we investigated whether the performance of the HNN depended on the brain regions of the recorded neurons. We observed improvements in HNN performance in almost all the recorded brain regions except the HPC (Wilcoxon signed rank test: AC, PPC, STR, M1, and OFC, p = 0.030–6.2e-6; HPC p = 0.080) (**Fig. 2c)**. We also examined whether the performance of the HNN depended on the number of recorded neurons. To normalize the variability in RMSE across sessions, we used RMSE improvement ratios (i.e., the proportions of RMSEs in the HNN that changed from those in the ANN). We found that the RMSE improvement ratios were correlated with the number of neurons (**Fig. 2e**) (**Methods**). When analyzed by brain region, the number of neurons in the PPC and OFC showed significant correlations (**Fig. 2e**). To explore this further, we selected sessions containing data from at least 40 neurons and investigated how the RMSE changed when we randomly selected 1, 5, 10, 20, or 40 neurons from the same neuron group (**Fig. 2f**). This method reduced the variabilities in HNN performance caused by the sessions (i.e., differences in recorded neurons and mouse body movements). We found that the HNN performance depended on the number of spike inputs, especially from the PPC (**Fig. 2f**). We then directly compared the RMSE improvement ratios across brain regions and found that using PPC neurons yielded slightly better predictions of body movements than neurons from other regions (**Fig. 2g**). These results suggest that the PPC had distributed representations of body movements across neurons.

### Compared with the artificial units in ANNs, those in HNNs exhibit more neuron-like activity

To investigate how the HNN achieved better performance than the ANN, we first examined the output weights that connect each unit to body movement. When comparing the mean absolute output weights per unit across models, we found that HNN units had significantly smaller output weights than those in ANN (**Fig. 3a**), consistent with a previous finding showing that small output weights are associated with better generalization and reduced overfitting^25^, consistent with the performance in HNN (**Fig. 2**).

**Figure 3.**
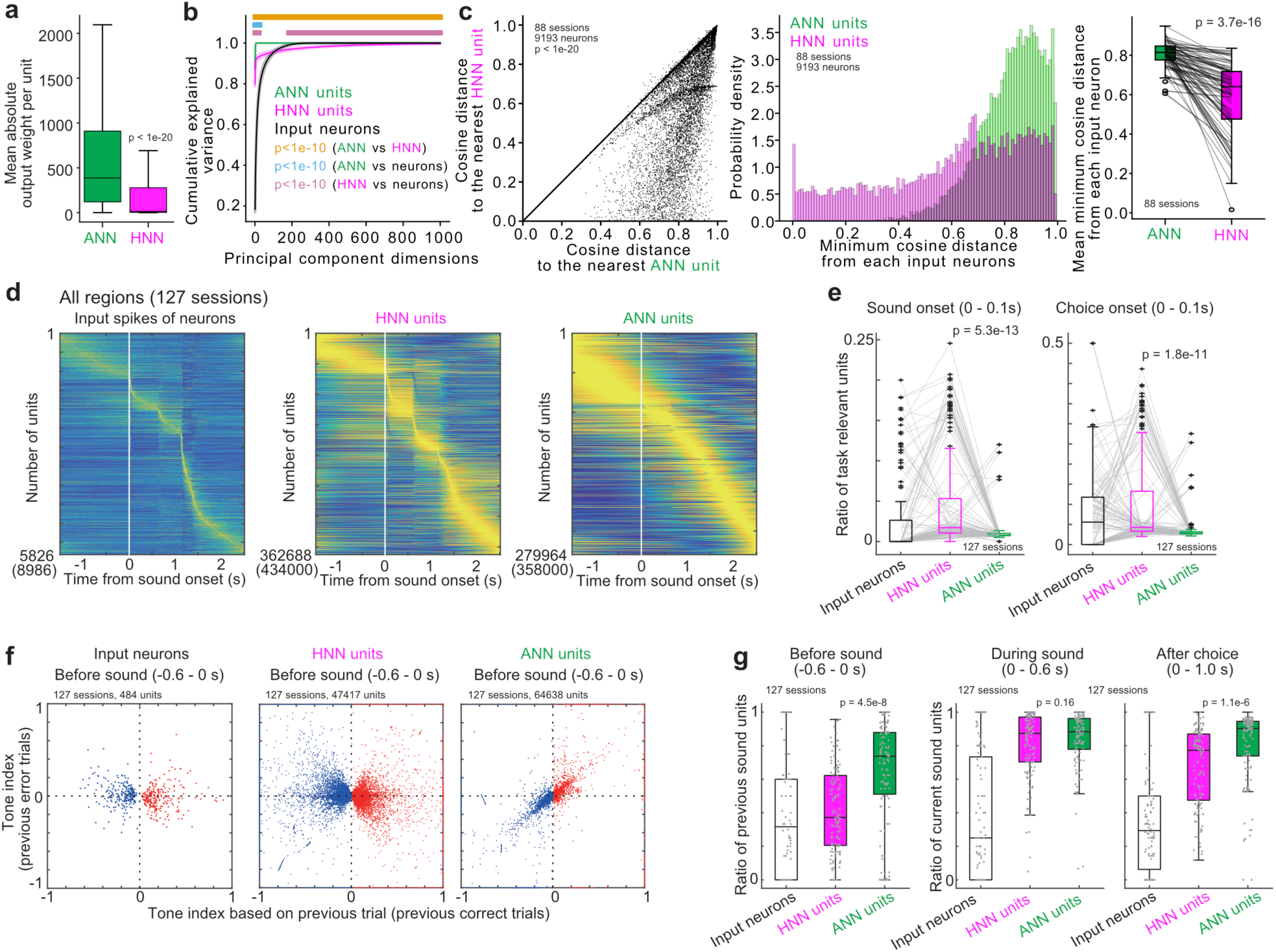
The activity of artificial units in the HNN is similar to that of neurons in mice. **a.** Output weight of each unit in ANN and HNN (190,449 and 316,358 units, respectively). The output weights of HNN units were significantly smaller than those of ANN (Mann‒ Whitney U test). Boxplots are the same as those in Fig. 2d. **b.** Cumulative explained variance from principal component analysis for ANN or HNN units or input neurons. Means and standard errors of mean (88 sessions from 6 mice for the ANN, HNN units and input neurons). When the dimensions of input neurons (median 51.5) beyond the number of neurons, we set the explained variance as 1. For the dimensions beyond 184, the explained variance of input neurons was consistently lower than that of the HNN units (Wilcoxon signed-rank test). **c.** Cosine distances from each mouse neural activity to the nearest ANN or HNN unit activity. Each neuron’s cosine distance to its nearest HNN and ANN unit were scatter-plotted (left). The distribution of minimum cosine distances is shown as a histogram (Center). For each session, the average of the minimum cosine distance was calculated for ANN and HNN units (Right). The activity of neurons in mice was significantly closer to that of units in HNN than to that of ANN (Wilcoxon signed-rank test). Boxplots are the same as those in **a**. **d.** Average activity of task-relevant units with increased activity during the task. The spike input of neurons is presented as the average activity of mouse neurons across trials from the AC, PPC, HPC, STR, M1, and OFC. The HNN and ANN units are presented as the average activities of the artificial units. Parentheses show the number of all units across sessions. The order of task-relevant units was sorted according to the maximum activity timings. The average activity of each unit was standardized between 0 and 1. **e.** Proportion of task-relevant units at sound and choice onsets in each session. Compared with those of the ANN, the activities of the artificial units in the HNN were aligned with the task events (Wilcoxon signed-rank test). Boxplots are the same as those in **a**. **f.** Tone indices of mouse neurons and artificial units before sound. We analyzed the task-relevant units with increased activity before the sound (-0.6–0 s) and significant differences in activity between the previous correct trials with low- and high-category tones (p < 0.01 in the Mann‒Whitney U test). The tone indices of the mouse neurons and HNN units were widely distributed. **g.** Ratio between the sound- and choice-responsive units. The sound- and choice-responsive units had tone indices with the same and opposite signs in the correct and incorrect trials, respectively. Similar to **c**, we analyzed the task-relevant units with increased activity at the corresponding time windows and different activities between the low- and high-category correct trials. The ratio of sound-responsive neurons was smaller in the HNN units than in the ANN units before sound and after choice (Wilcoxon signed-rank test). Boxplots are the same as those in **a**.

We analyzed the internal dynamics of the artificial units of RC (**Fig. 3b-d**). As the number of HNN units was larger than the input neurons and the HNN received more inputs than the ANN, the HNN units required a larger number of principal components to achieve high cumulative explained variance (**Fig. 3b**). We then used cosine distance to quantify the similarity between the activities of real neurons and those of HNN and ANN units. The results showed that neuronal activity was more similar to the HNN units than to the ANN units (**Fig. 3c**).

Among all the neurons recorded from the AC, PPC, HPC, STR, M1, and OFC (**Fig. 2e**), we investigated the task-relevant neurons that exhibited increased activity during the task, as in our previous studies^7,21,26^ (**Methods**). Specifically, task-relevant neurons showed increased activity in at least one time window (0.1 s) between -1.5 s and 2.5 s from the sound onset (40 windows) or between -0.5 s and 2.5 s from the choice (30 windows, 40 + 30 = 70 windows in total) compared with the baseline activity (p < 1e-10 in one-sided Wilcoxon signed-rank test). The baseline activity was recorded from -0.2 s to 0 s from spout removal and -0.2 s to 0 s from spout approach for the sound- and choice-aligned activities, respectively. Using the same criteria, we identified the task-relevant artificial units in the HNN and ANN based on the activity of the RC units. We found that the activity of neurons in mice and that of artificial units in the HNN were aligned with the task events of sounds or choices (**Fig. 3d**). We analyzed the proportion of task-relevant neurons activated at sound and choice onsets and verified that the activity of HNN units was more closely related to the task events than was that of the ANN (**Fig. 3e**).

In addition to analyzing the temporal patterns of artificial units, we analyzed the representations of real neurons and artificial units by introducing the tone index (**Methods**)^7,21^. The tone index was used to compare the activity during the low- and high-category tone trials and ranged between -1 and 1. We independently analyzed the tone indices for correct and incorrect trials based on the neural activity before the sound, during the sound, and after the choice. When neurons had significantly different activities during correct low- and high-category tone trials (p < 0.01 in the Mann‒Whitney U test) and the tone indices of correct and incorrect trials had the same signs, we defined the neurons as sound responsive. When the signs of the tone indices of correct and incorrect trials were opposite, we defined the neurons as choice responsive.

We analyzed the tone indices before sound according to the tone categories in previous trials (**Fig. 3f**). We found that the mouse neurons and the HNN units had distributed tone indices, whereas the ANN units had the same signs of tone indices between the correct and incorrect trials, suggesting that the ANN units represented previous sound categories, which was inconsistent with the neurons in mice. We compared the ratio of sound-responsive to choice-responsive neurons in three different time windows in each session (**Fig. 3g**). We found that a greater proportion of artificial units in the HNN were choice responsive than the ANN before sound and after choice, which is consistent with the findings in neurons in mice.

### The population trajectories of artificial units in HNN exhibit choice modulations before choice onset, which is consistent with mouse neurons

Decoding analyses using the generated body movements from the HNN and ANN revealed that the HNN achieved faster choice separations than the ANN, consistent with the findings in mice (**Fig. 1e**). Single-unit analyses further indicated that the artificial units of the HNN represented the choices (**Fig. 3**). To further investigate choice representations in the HNN, we investigated the population trajectories of HNN units and compared them with those of mouse neurons and ANN units (**Fig. 4**). For this analysis, we used 97 sessions out of 127 sessions in which at least 20 neurons were recorded in each brain region. We found that the population decoding trajectories were diverged between left and right choices at -0.1–0 s in the mouse neurons, HNN units and ANN units (Wilcoxon signed-rank test: p = 4.0e-3–4.3e-12) (**Fig. 4a-c**). Direct comparisons of choice separations revealed that the HNN units had significantly greater separations than the mouse neurons and ANN units did (**Fig. 4d, e**).

**Figure 4.**
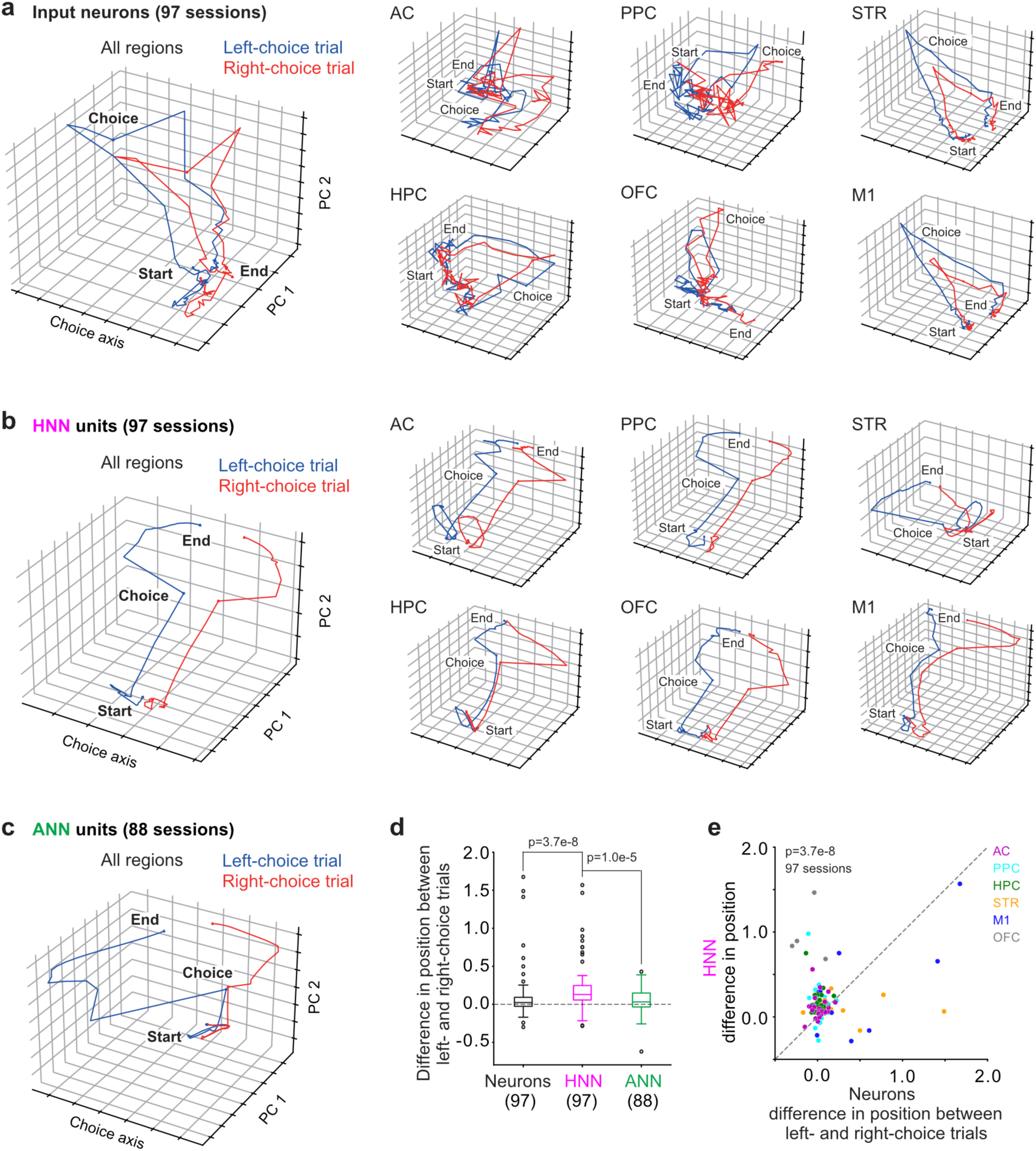
Choice separations of population trajectories in artificial units of the HNN. **a-c.** Population trajectories of choice separations. Average population trajectories of the neurons in the mice (**a**), the artificial units of the HNN (**b**), and the artificial units of the ANN (**c**) are shown. For the mouse neurons and HNN units, the average trajectory in each brain region is also shown. **d.** Comparisons of choice separations at -0.1–0 s from choices. The HNN units had larger separations of choices than did the mouse neurons and ANN units (Mann‒Whitney U test). Boxplots are the same as those in Fig. 2d. **e.** Comparisons of choice separations between the HNN units and mouse neurons. The data are the same as **d**, but colored according to the brain regions of neurons.

These results suggest that the HNN generates choice-relevant unit activity by utilizing, propagating, and amplifying the partially recorded neural activity. Together with the result that the behavioral prediction accuracy in the ANNSP was lower than that in the HNN (**Fig. 2b**), this indicates that neural-activity inputs were integrated with task parameters and processed within the HNN, thereby generating additional information relevant to behavior.

### Artificial units in the HNN compensate for the unrecorded neural activity of mice

A potential advantage of the HNN is that the artificial units can predict the neural activity of unrecorded neurons for predicting how the whole brain generates body movements for decision-making. To test this hypothesis, we first examined whether the HNN could generate the activity patterns corresponding to unrecorded neurons by quantifying the similarity between unit activity and neuronal activity using cosine distance.

We selected mouse neurons that were close to ANN units based on cosine distance, defined as near neurons, and constructed a “near-HNN” using only the near neurons as the inputs of RC. We then compared the cosine distances across four categories: (1) the HNN units, (2) the ANN units, (3) near neurons, and (4) far neurons that were not used as the inputs of near-HNN. The threshold of cosine distances between the near and far neurons were heuristically determined as 0.95. We found that the activity of near-HNN units was more similar to that of the far neurons than that of ANN units, both at the single-neuron level (**Fig. 5a**) and across sessions (**Fig. 5b**). These results suggest that the HNN generated unit activity that was similar to the mouse neuronal activity, compared to ANN, regardless of whether those neurons were used as the inputs of RC.

**Figure 5.**
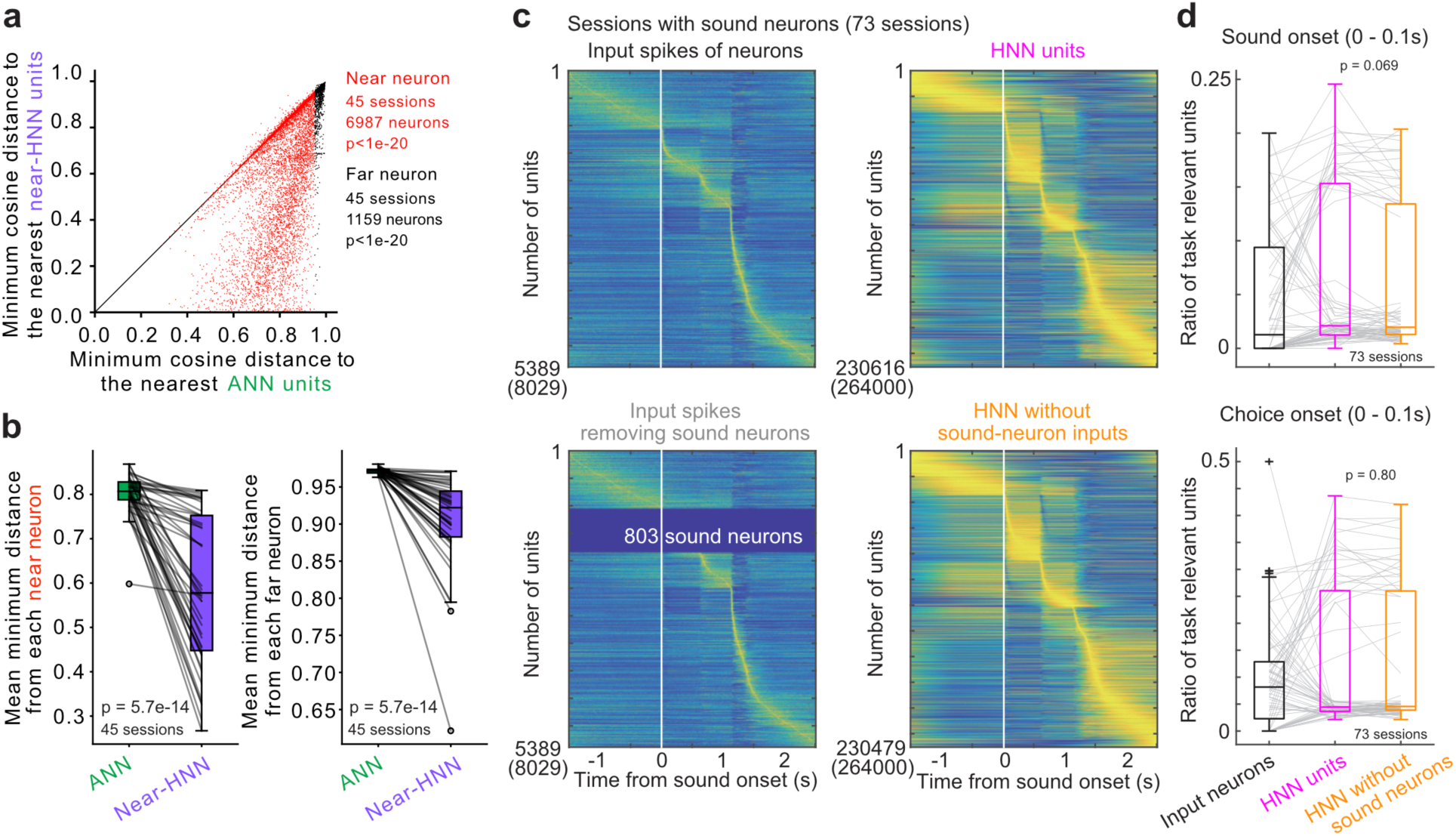
Artificial units in the HNN compensate for unrecorded neural activity of mice. **a.** Cosine distances from each mouse neural activity to the nearest ANN or HNN unit activity. Input neurons were categorized as far or near neurons according to the cosine distance from ANN units (threshold, 0.95). We selected 45 sessions which had at least five far neurons to avoid bias from single-neuron variabilities. The activity of mouse neurons was significantly more similar to that of HNN units than to that of ANN units (Wilcoxon signed-rank test). **b.** Mean of minimum cosine distance between each mouse neuron and the nearest unit in either the ANN or HNN for each session. The average of the minimum cosine distances was calculated separately for the ANN and HNN. The average cosine distances of near and far neurons were smaller in near-HNN units than in ANN units (Wilcoxon signed-rank test). Boxplots are the same as those in Fig. 2d. **c.** Average activity of task-relevant units with increased activity during the task. The data plots are the same as those in Fig. 3d, but we used the 73 sessions in which the sound neurons were recorded. HNNs using spike inputs with (top) and without (bottom) sound neurons are shown. **d.** Proportion of task-relevant units at sound and choice onsets in each session. The spike inputs with or without the sound neurons did not significantly affect the proportion of sound- and choice-onset activity of artificial units in the HNN (Wilcoxon signed-rank test). The data plots are the same as those in Fig. 3e but from 73 sessions.

To further investigate whether the HNN compensated for the activity of unrecorded neurons, we examined the activation timing of HNN units around sound onset. We removed the task-relevant neurons that exhibited increased activity during sound presentation (0.6 s) (defined as sound neurons) from the inputs of the HNN and trained the RC without these neurons (**Fig. 5c**). In total, we removed 803 out of 5389 task-relevant neurons, which were recorded in 73 out of 127 sessions. We then compared the activity of HNN artificial units with and without the inputs of sound neurons across these 73 sessions (**Fig. 5d**). Even in the absence of input from the sound neurons, the artificial units in the HNN remained active at sound and choice onset. These results suggest that the HNN generates neuron-like activity patterns by compensating for missing inputs through artificial units in recurrent neural network dynamics.

## Discussion

In this study, we developed a hybrid neural network (HNN) model that uses the activity of real neurons in mice as inputs for reservoir computing (RC). Our model predicted the body movements of mice using the sensory inputs of sounds, rewards, and spout movements. The sensory inputs were presented to the mice during the tone-frequency discrimination task in our previous study^26^.

We found that the performance of the HNN was better than that of an artificial neural network (ANN) with only artificial units (**Fig. 2**). The RC with spike inputs alone did not outperform the RC with only the task inputs (ANN), suggesting that the recorded spikes did not capture all the sensory inputs. The HNN was independently developed using neurons from one or two brain regions in each session. To capture all the sensory inputs, further experiments that simultaneously record 1) the neural activity across multiple brain regions and 2) a large population of neurons with a wide-field two-photon microscope^27–29^ or multiple Neuropixels probes^30^ are needed.

We found that the performance of RC was improved by spike inputs from almost all the recorded regions except the HPC (**Fig. 2c**). As previous studies have shown that neural activity is modulated by body and facial movements^31–34^, it was not surprising that the neurons in many brain regions contributed to generating movements on a continuous time scale. The HPC may not be directly involved in controlling body movements, but is instead considered to play a role in higher-order aspects of behavioral control, such as action planning, sequence prediction, and prospective coding of future behavior^35–37^. We also found that the accuracy of the HNN was improved by increasing the number of spike inputs from the PPC (**Fig. 2f**). The PPC not only represents choices in perceptual decision making^38^ but also changes activity during inter-trial intervals^39^ and estimates stimulus probabilities^40^, expected outcomes^41,42^, and hidden contexts^43^. Assuming that such diverse internal representations are distributed across multiple neurons, the accuracy of body movement prediction may increase in a neuron number-dependent manner. Although M1 is thought to be the center of behavior output and its improvement was statistically significant, the effect was relatively moderate compared to other areas. Previous studies suggest that the primary motor cortex (M1) is deeply involved in action initiation, controlling movement direction and force parameters, and adaptive fine-tuning. On the other hand, there is also evidence indicating that M1 does not faithfully map the spatial coordinates of each body part^44–46^. Thus, for a task like ours that requires predicting the coordinates of body parts, relying on M1 activity may not have yielded sufficient performance.

Next, we found that the activity of HNN units was more similar to spike activity than that of the ANN (**Fig. 3**). In addition, the HNN outperformed the ANNSP in behavior prediction (**Fig. 2b**). PCA showed that the HNN representations were higher-dimensional than the original neuronal activity (**Fig. 3b**). Furthermore, population decoding analysis showed that the HNN achieved better choice separation than both the ANN and real neurons (**Fig. 4**). Taken together, these results suggest that the activity of HNN units contains movement information beyond that present in the original neural activity.

We investigated whether the artificial units of our HNN could predict the neural activity of unrecorded neurons (**Fig. 5**). Even when we developed the HNN without the inputs of neurons with specific activity pattern, the HNN units still exhibited activity patterns that resembled those of the omitted neurons, suggesting that the HNN could compensated for the missing neural activity. In addition, even when sound-relevant neurons were excluded from the input, the HNN units were activated at the time of sound presentation. Notably, the generated HNN units’ activity accurately predicted behavior with significantly smaller output weights than the ANN. Instead, its internal unit dynamics were sufficient to predict body movements. To further investigate the functional role of these internally generated unit activities and whether they compensated for neural activity in specific areas, future work could utilize simultaneous recordings from multiple brain areas. Such data would allow us to assess whether inputs from one area can generate activity in another area.

We developed the HNN using a simple reservoir computing architecture in which the internal weights were not updated from their initial random values by assuming that the brain itself utilizes somewhat random connectivity to propagate information^47^. Although this simple RC architecture led to notable improvements of behavioral prediction, as shown in this study, it is likely that further enhancements could be achieved by incorporating architectures with trainable internal connections.

One advantage of developing brain-like AI is that pseudoinactivation of the HNN may simulate changes in mouse behavior in inactivation experiments using optogenetics or other methods (**Fig. S4**) (**Supplementary Methods**). To test this hypothesis, we trained an HNN using the neural activity of the OFC as the input, as in **Fig. 1**. We then removed the spike inputs from the HNN to represent pseudoinactivation (**Fig. S4**). We found that although optogenetic inhibition of the OFC significantly reduced tone history-dependent choice biases in mice during the task, pseudoinactivation of the HNN only slightly reduced discrete choice biases. This discrepancy may be explained by the relatively small number of recorded OFC neurons used as input to the HNN; pseudo-inactivating only a small subset of OFC neurons may have a limited effect compared to the broad suppression achieved by optogenetics. These findings suggest that accurately replicating the effects of optogenetic inactivation may require recording from a larger population of OFC neurons, or, beyond merely silencing recorded inputs, targeting HNN units that exhibit OFC-like dynamics.

In summary, we developed an HNN that integrates real neurons from mice with artificial units to achieve more accurate predictions of mouse body movements than a conventional ANN composed solely of artificial units. The activity of the HNN’s artificial units was more similar to that of real neurons than that of the ANN, as measured by cosine distance. These units were temporally aligned with task events and reflected the mice’s choices—mirroring the activity patterns of actual neurons. Remarkably, simply inputting neural activity into a reservoir computing network—without updating internal weights—enabled the HNN to compensate for unrecorded neural activity. While further validation using a larger number of spike inputs and alternative architectures will be essential, this HNN offers a promising approach for understanding brain function by data augmentation: propagating observed neural activity through a randomly connected network to extend information content and improve behavior prediction accuracy.

## Methods

We developed a real-cyber hybrid neural network (HNN) in which the neural activity of mice during a behavioral task was used as the input for reservoir computing (RC). The animal experiments and datasets used in this study were reported in detail in our previous study^26^.

### Animal experiments

All animal procedures were approved by the Animal Care and Use Committee at the Institute for Quantitative Biosciences (IQB), The University of Tokyo. We used 14 male CBA/J mice (Charles River, Japan) that were 8 to 15 weeks of age at the start of behavioral training with electrophysiology^26^. We recorded the body movements of 6 of the 14 mice with either 4 or 5 cameras (1 and 5 mice, respectively) placed around each mouse during a task. The data from these 6 mice were used in this study. The mice were housed in a temperature-controlled room with a 12 h/12 h light/dark cycle. Before surgery, 3 mice were housed in one cage. The mice were allowed ad libitum access to food, and their water intake was restricted to 1.5 mL per day. The weights of the mice were checked daily to prevent dehydration. The mice were caged in isolation after craniotomy.

The surgical procedures used were previously described^7,21,26,48^. Our surgery involved two steps: the implantation of a lightweight head bar and a craniotomy for electrophysiology. In each surgery, the mice were anesthetized via an intraperitoneal injection of a mixture of medetomidine (0.3 mg/kg), midazolam (4.0 mg/kg), and butorphanol (5.0 mg/kg) or isoflurane (2% at induction, below 1.5% for maintenance). Meloxicam (2 mg/kg, subcutaneous) and eye ointment were also used. The electrophysiology recording site was chosen from the following candidates: the recording sites of the OFC were centered at +2.6 mm anterior–posterior (AP) and ±1.4 mm medio–lateral (ML) from bregma; the sites for the STR and M1 were +0.8 mm AP and ±1.5 mm ML; the sites for the PPC and HPC were -2.0 mm AP and ±1.7 mm ML; and the sites for the AC were -3.0 mm AP and ± 3.8 mm ML.

#### Electrophysiology during a tone-frequency discrimination task

The behavioral task was explained in detail in our previous study^26^. Briefly, the task was based on the tone-frequency discrimination task of previous studies^22,23,49^. The mice were head fixed and positioned over a cylinder treadmill. One precalibrated speaker (#60108, Avisoft Bioacoustics) was placed diagonally to the right of each mouse for auditory stimulation (Type 4939, Brüel and Kjaer). Water was delivered through two spouts connected to an infrared lick sensor (Ohara, Inc.). The behavioral system was controlled by a custom MATLAB (MathWorks) program running on the Bpod r0.5 framework (https://sanworks.io) in Windows.

Each mouse selected the left or right spout depending on the dominant frequency of a tone cloud to receive a water reward^22,49^. The frequency of each sound pulse in the tone cloud was sampled from 18 logarithmically spaced slots (5-40 kHz). The tone cloud in each trial contained low-frequency (5–10 kHz) and high-frequency (20–40 kHz) sounds and was categorized as low or high depending on the dominant frequency. The high- or low-category tone in a correct trial was associated with a reward of 2.4 μl of 10% sucrose water in the left or right spout. An incorrect trial resulted in a noise sound burst for 0.2 s. When the mice did not select a spout within 15 s from the start of the trial, a new trial started.

Our task had two task conditions for sound presentations on the basis of the transition probability ‘p’ of the tone category in each trial^26^. In the repeating condition, the tone category was repeated often in each trial (p = 0.2), whereas in the alternating condition, the tone category was altered by 80% (p = 0.8). Each mouse was assigned to one of the two task conditions (1 and 5 mice assigned to the repeating and alternating conditions, respectively).

All the electrophysiological recordings were performed with Neuropixels 1.0 (IMEC) inside the sound-attenuating booth (Ohara, Inc.). A subset of neurons from 6 out of 14 mice reported in our previous study were used^26^. Every day, i.e., every session of the behavioral task, we inserted one Neuropixels probe that targeted either the OFC, STR/M1, PPC/HPC or AC. The probe location was identified in the post hoc fixed brain by labeling the probe injection site with CM-DiI or DiA at the recording phase (Thermo Fisher Product #V22888 or #D3883). The Open-Ephys GUI was used to acquire the neural data at a sampling rate of 30 kHz with a gain of 500 (PXIe acquisition module, IMEC)^50^. The task events were sampled at 2.5 kHz (BNC-2110, National Instruments). Spike sorting and manual curation were performed with Kilosort 3 on MATLAB (https://github.com/cortex-lab/KiloSort) and Phy on Python (https://github.com/cortex-lab/phy).

We defined the depth of each unit according to the approximate location of the probe and the electrode position, which measured the average maximum amplitude of spikes. For the OFC recording, the units with an estimated spike depth from the brain surface less than 1.9 mm were analyzed. For the PPC or HPC, the units with an estimated spike depth less than 1.0 mm or between 1 mm and 2.3 mm, respectively, were analyzed. For the STR or M1, the units with an estimated spike depth greater than or less than 1.5 mm, respectively, were analyzed. All the recorded units for the AC probe were analyzed.

As described in our previous studies^7,26^, after electrophysiological recording, the mice were deeply anesthetized with isoflurane (5%) and further anesthetized with a mixture of 1.5 mg/kg medetomidine, 20 mg/kg midazolam, and 25 mg/kg butorphanol. The mice were perfused with 10% formalin solution to identify the probe locations on fixed brain slices.

### Measurement of mouse body movement

During the tone-frequency discrimination task, we captured the mouse body movement with 5 video cameras in 5 mice across 69 sessions (DMK 33UX273, Imaging Source). For the other mouse, we captured body movements with 4 cameras in the first 8 sessions and 5 cameras for the remaining 11 sessions (88 sessions in total) (ICCapture Express). The front camera was placed in front of the face of the mouse to capture tongue movement. The front left and front right cameras were placed at the side of the mouse’s face to capture the forelimbs and whiskers. The other 1 or 2 back cameras, depending on the setting, were placed behind the mouse to capture the trunk, hind limbs, and tail. The frame rate of the video was 137 Hz on average, with a standard deviation of 0.00046 Hz, for the first 53 sessions and 140 Hz, with a standard deviation of 0.00237 Hz, for the last 35 sessions. The frame rates were different both without and with the light-emitting-diode (LED) devices used for video alignment.

From the video data from each camera, we first extracted the XY trajectories of body points with DeepLabCut (DLC)^24^. From the front camera data, we extracted 6 body points, namely, nose, tongue, right foreleg finger, right foreleg wrist, left foreleg finger, and left foreleg wrist. From the front left and front right camera data, we extracted 7 positions each, namely, nose, side upper eye, side bottom eye, right foreleg finger, right foreleg wrist, left foreleg finger, and left foreleg wrist. From the back camera data, we extracted 10 positions each, namely, right hindleg toe, right hindleg heel, right hindleg knee, left hindleg toe, left hindleg heel, left hindleg knee, back top, tail tip, tail middle, and tail root. In total, we extracted 30 and 40 body points from the 4- and 5-camera settings, respectively. In addition to the body points, we also extracted the movements of the spouts in each of the videos for temporal alignment.

We used data from several sessions to train the DLC: 10 sessions (2 sessions each from 5 mice) for the front camera, 5 sessions (1 session each from 5 mice) for the front left and front right cameras, 1 session from 1 mouse for the back camera, and 5 sessions (1 session each from 5 mice) for the back left and back right cameras. In each session, we randomly sampled approximately 100 frames from each video and manually labeled the body points. We extracted 996, 500, 500, 100, 500, and 499 frames for the front, front left, front right, back, back left, and back right cameras, respectively. The labeled frames from the same camera were used to train one DLC model, which was applied to all the sessions from the corresponding camera across the mice. After the automatic extraction of body points with DLC, we detected outlier points with a likelihood of less than 0.5. These outliers were replaced with linear interpolations with the nearest points with likelihood above 0.5. The trajectory of each body point was z scored with a mean of 0 and a standard deviation of 1.

For the back cameras, the appearance of spout movements differed across sessions. We thus trained 2 and 3 DLC models for the back and back left cameras, respectively, and applied the proper DLC model for spout movement extraction.

### Time alignment of movie data from all the cameras and task parameters

To temporally align the movies from different cameras, we extracted the spout movements in all the movies with DLC in 55 out of 88 sessions in 6 mice. In the other 33 sessions, all the cameras detected a 10-ms infrared light stimulus (660 nm) from the LED in each trial, which was used for time alignment. The LED stimulus was also sampled at 2.5 kHz as the task parameter (BNC-2110, National Instruments).

#### Time alignment with spout movements

For each camera, we detected the spout movement by using either the x or y coordinate with a large variance. On the basis of the spout coordinates from all the cameras, we used the *finddelay* function in MATLAB to analyze the differences in movie frames among cameras. We then delete the initial and final frames in each camera to coarsely align the camera data. The aligned spout coordinates from all the cameras were combined and averaged. Using the spout movement detected by a rotary encoder (BNC-2110) with a sampling rate of 2.5 kHz and the average spout movement from all the cameras, we determined the frames per second (fps) of the video data. The average fps was 140 Hz (35 sessions) or 137 Hz (53 sessions). We then downsampled the task events that were originally sampled at 2.5 kHz to match the camera fps, ensuring that no task events were lost in the process.

Subsequently, fine discrepancies due to frame drops or other issues in the behavioral data from each camera were corrected on a trial-by-trial basis. For the spout data obtained from the input data and from 4 or 5 cameras, we calculated a moving median over 201 frames (100 frames before and after). In parallel, thresholds were calculated by applying a moving average over 30,001 frames for all of the spout data. The data were then binarized to 0 or 1 depending on whether the spout moving medians were above or below each threshold. The frame numbers at which the values of these binarized spout data changed were identified as the spout movement timing frame numbers for each trial. By calculating the difference in spout movement frame numbers between the input data and the camera data for each trial, the error in frame count for the spout movement timing was obtained for each trial.

A moving median over 21 trials (10 trials before and after) was calculated for the trial-by-trial spout movement timing error data. The moving median was used to avoid overcorrecting for trial-specific noise influenced by errors in the automatic labeling by DLC. Finally, the body coordinate data were adjusted by reducing or increasing the number of frames by the amount of this moving-median error frame count, followed by linear interpolation. This linear interpolation was applied to all body coordinate data, including the spout movement data, thereby temporally aligning the coordinate data with the input data.

#### Time alignment with LED

In each video, we analyzed the average luminance of a rectangular region including the LED stimulus in each frame. By computing the differences between the luminance values in consecutive frames and setting any negative differences to zero, we created LED data that contained positive values for the frames where the LED was lit. We calculated the moving median for these LED data over a total of 301 frames (150 frames before and after) and subtracted it from the LED data to eliminate luminance changes with periods longer than 300 frames. The corrected LED data were then z scored (i.e., mean 0, std 1). A threshold was manually determined (either 0.3 or 0.2), and the data were binarized to 0 or 1 according to this threshold. In sessions where incorrect threshold crossings were visually confirmed, we set the value to 1 only when the threshold-crossing frame was at least 500 frames after the previous frame, since intervals between trials were always more than 500 frames. The frame numbers at the moments when the value changed from 0 to 1 (LED-on frame numbers) were obtained and used for time alignment.

First, to align the frame numbers of the first LED-on frames across the five camera images, we deleted the initial frames from the body coordinate data of the five cameras so that the first LED-on frames aligned. Next, for each of the body coordinate datasets 1 to 5, we obtained the LED-on frame numbers of the first and last trials and calculated the total number of frames between them, i.e., the total frame count for all trials. Similarly, we obtained the frame numbers corresponding to the trial initiation timings in the input data (which are the same as the LED emission timings) and calculated the total frame count for all trials between the first and last trials.

By comparing the longest total frame count among the 5 body datasets with that of the 1000 Hz input data, we determined the fps of the body data. The input data were then uniformly downsampled by deleting frames at regular intervals to match this fps.

Finally, to ensure that the number of frames per trial was consistent, we performed linear interpolation so that the LED-on timings in body datasets 1 to 5 aligned with the trial initiation timings in the input data for each trial. During interpolation, we recorded the number of frames deleted or added for each trial and manually confirmed that there were no obviously incorrect frame adjustments exceeding 100 frames.

### Data analysis

Artificial neural networks were programmed with Python (3.11.4 and 3.6.10). The data were analyzed with MATLAB (MathWorks) and Python (3.11.4 and 3.6.10).

We used the data from 6 out of 14 mice from our previous study^26^, as body movements were captured by cameras only in these 6 mice over 88 sessions. The neural data from the PPC-HPC and STR-M1 were simultaneously recorded with one Neuropixels probe. We analyzed the data from 27 sessions for the PPC-HPC (no neurons in the PPC in 2 sessions or the HPC in 1 session), 15 sessions for the STR-M1, 26 sessions for the AC, and 20 sessions for the OFC. Mice u01 – u06 had data from 19, 17, 20, 2, 19, and 11 sessions, respectively. We selected the sessions for analyses using the same criteria used in our previous study^26^: (i) the percentage of correct responses for both the 100% low-tone and 100% high-tone stimuli was greater than 75%, and (ii) the total reward amount in one session was at least 600 µL.

### Modeling the body movements of the mice

#### Artificial neural network (ANN) with reservoir computing

We used reservoir computing (RC) (ReservoirPy, Python), a type of recurrent neural network^12,17^, to generate the body movements of the mice during the tone-frequency discrimination task. The RC was trained on the weights of the full connections between the recurrent units and outputs via ridge regression. The recurrent units were leaky-integrator neurons with a leaking rate *lr* of either 0.05 or 1. The connections (i) between the RC inputs and the recurrent units and (ii) between different recurrent units were probabilistically determined, and the probability was optimized as the hyperparameters, described later. The activation function of the recurrent units was the hyperbolic tangent (*tanh*) function. The update of states was described with the following equation with *x*_!_ representing the current state, *x*_!“#_representing the next state, *u* representing the input vector, *W*_$%_representing the weight between inputs and RC units, and *W* representing weights among RC units:

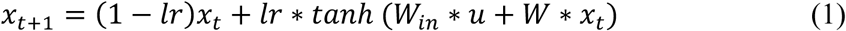

The RC inputs were the short pure-tone pulses in the tone cloud (low or high category), left or right water reward, error tone, and spout movement, which were downsampled to 140 Hz (35 sessions) or 137 Hz (53 sessions) to align with the movie data. The water rewards were defined to deliver between 0 and 20 ms from the reward onset. All the inputs ranged between 0 and 1. There were 6 task inputs in total (**Fig. 1a**).

#### Real-cyber hybrid neural network (HNN)

Our HNN used recorded neuronal activity as the RC inputs as well as the 6 task inputs. The neuronal activity was averaged and downsampled to 140 Hz (35 sessions) or 137 Hz (53 sessions). The spike counts in each time window were scaled and ranged between 0 and 1.

In addition to using time-aligned mouse neural activity as the RC inputs, we used time-shifted activity as the input to investigate whether the precise timing of neural activity contributed to generating the body movements of the mice (**Fig. 2c**). We used the neural activity before 3 s as the input of the HNN by putting the last 3-s activity at the initial timing of the session. We also constructed the HNN with spike inputs but without the task inputs.

#### Training and testing of the ANN and HNN

In each session, we used the first 80% of the data for training the RC and the last 20% of the data for testing. Within the training data, we optimized the hyperparameters (**Fig. S1**). We tested the 2 leaking rates of each unit (0.05 or 1) and the 44 combinations of average connections per unit and number of nodes (for 1 or 3 connections per unit: 1000 to 7000 nodes with every 1000 increments; for 2, 4, or 5 connections: 1000 to 5000 nodes with every 1000 nodes; for 10, 30, or 100 connections: 3000 to 7000 nodes with every 1000 nodes). There were 88 hyperparameters in total. The spectral radius (*sr* = 0.9), input scaling (*iss* = 1), and ridge penalty (*ridge* = 1e-7) were fixed hyperparameters.

We used the first 80% of the training data (the first 64% of all the data in each session) and trained the RC with each of the 88 hyperparameters. The root mean square error (RMSE) of the RC was determined with the last 20% of the training data (16% of all the data) to identify the hyperparameter setting with the smallest RMSE. The RMSE was defined as the average RMSE of all the XY trajectories of body points (60 or 80 XY trajectories for 4 or 5 camera settings, respectively). Using the identified hyperparameter setting, we trained the RC with all the training data (80% of all the data) and analyzed the performance with the test data (the last 20% of all the data); the test data were used in neither the hyperparameter setting nor the connection weights of the RC.

We defined the RMSE improvement ratio of the HNN on the basis of the RMSE with no spike inputs (i.e., ANN) to evaluate the relative increase in performance by adding spike inputs (**Fig. 2e, g, S3b**):

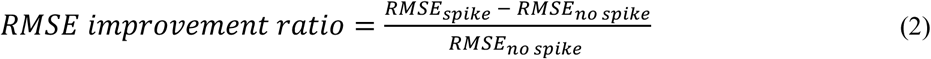

When we analyzed the performance of HNN with 1, 5, 10, 20, or 40 input spikes (**Fig. 2f, g**), we used the hyperparameter settings for which the RC achieved the best median performances in 23 sessions of 1–5 spike inputs (1 and 5 neurons), 27 sessions of 6–15 spike inputs (10 neurons), 20 sessions of 16–29 spike inputs (20 neurons), and 23 sessions of 30–50 spike inputs (40 neurons) (**Fig. S1**). In each session, we randomly selected neurons with spike inputs of 1, 5, 10, 20, and 40 for 10 times and performed HNN to analyze the median RMSE for the 10 iterations.

#### Average model

We divided each trial into four sections: (1) before tone onset, (2) between tone onset and spout arrival, (3) between spout arrival and choice, and (4) after choice. In each section, we divided the trials into different categories and analyzed the mean trajectories of body points: Section 1 had one category; Section 2 had low- and high-tone categories; Section 3 had one category; and Section 4 had three categories: left reward, right reward and error tone (**Fig. 2a**). The mean trajectories of the body points were analyzed using the first 80% of the data in each session (training data). We then compared the trajectories of the last 20% of the data (test data) with those of the mean trajectory model to analyze the RMSE.

#### Generalized linear model

We used the MATLAB software package glmnet (*cvglmnet*) (https://glmnet.stanford.edu/index.html) (**Fig. 2a**)^51^. Our generalized linear model (GLM) included 60 kernels for each task event. The sound kernels (low and high) started from sound onset. The right reward, left reward, and error kernels started from the onset of the outcome. The spout kernels started from the arrival and departure of the spout. Each task-event kernel was 0.1 s and set for 6 s from the start. The initial 80% of the data in each session were used to train the GLM with 10-fold cross-validation (CV), whereas the last 20% of the data (test data) were used to compute the RMSE.

#### Decoding sound and choice categories from body movements

We decoded the sound categories (low or high) and the choices of the mice (left or right) from the body movements of the mice or those generated by RC (**Fig. 1e**). We used the MATLAB software package glmnet (https://glmnet.stanford.edu/index.html) with L1 regularization. In each time window of 0.1 s, we averaged the body movements and performed sound and choice decoding. The performance of the decoder was evaluated with 5-fold CV. Due to a shortage of test trials or computer memory, the 5-fold CV did not work in some sessions for the ANN and HNN; in these cases, we used the 3-fold CV. If the 3-fold CV did not work, we discarded the session. For decoding from the mouse data, we analyzed all the 88 sessions with 5-fold CV. For ANN, 62, 18, and 8 sessions were 5-fold CV, 3-fold CV, and discarded, respectively. For HNN, 65, 18, and 5 sessions were 5-fold CV, 3-fold CV, and discarded, respectively. To remove the contaminations of sound and choice decoding, we subsampled the trials used to train the decoder such that the number of trials for the 4 sound-choice combinations were identical (i.e., low-left, low-right, high-left, and high-right). We repeated the CVs 10 times with different subsamples and analyzed the average accuracy rates.

### Comparison of activity between neurons in mice and artificial units in RC

#### Cosine distance

To quantify the similarity between the activity of artificial units and real neurons, we compute the cosine distance between their activity vectors. The pairwise cosine distances were calculated as follows:

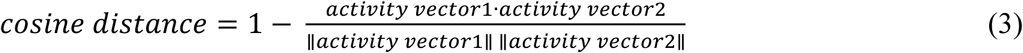

A smaller cosine distance indicates greater similarity in the temporal activity pattern.

#### Task relevant neurons and units

We analyzed the activity of neurons in mice and that of artificial units in the HNN and ANN on the basis of our previous studies^7,26^. We first identified the task-relevant neurons and units with increased activity during the task compared with baseline activity (one-sided Wilcoxon signed-rank test, p < 1e-10). Like in our previous study of mouse neurons^26^, for sound-aligned activity, task-relevant units presented increased activity in at least one time window (0.1 s) between -1.5 s and 2.5 s from sound onset (40 windows). For choice-aligned activity, task-relevant units exhibited increased activity in at least one time window between -0.5 and 2.5 s from the choice (30 windows, in total 40 + 30 = 70 windows). The baseline for the sound-aligned activity was -0.2 to 0 s from spout removal, i.e., before trial initiation. The baseline for the choice-aligned activity was -0.2 to 0 s from the time the spout approached, i.e., between the end of the sound and making a choice.

Using the average activity across trials, we identified the maximum activity timings of task-relevant units and compared the timings across units in the mice and models (**Fig. 3d**). We then analyzed the tone index of each unit on the basis of the left and right reward-associated tone trials^7,21,26^:

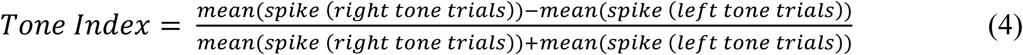

The tone index ranged between -1 and 1 and was independently analyzed for correct and incorrect trials. The tone indices before sound (-0.6–0 s) were analyzed on the basis of correct and incorrect previous trials, whereas the indices during sound (0–0.6 s) and after choice (0– 1.0 s) were analyzed for the current trials^26^. On the basis of the tone indices and the differences in activity between the left- and right-choice correct trials (p < 0.01 in the Mann‒Whitney U test), we analyzed the proportions of sound- and choice-responsive neurons before the sound, during the sound, and after the choice (**Fig. 3e, f**).

### Population decoding analysis of neurons in mice and artificial units in RC

We used a decoding-based analysis described in recent studies^52,53^. For the ANN, we analyzed all the 88 sessions. For the mouse neurons and HNN, we used 97 out of 127 sessions in which more than 20 neurons were simultaneously recorded from each brain region. All the population analyses were performed in each session separately (**Fig. 4**).

We first binned the data into 50 ms between -2.0 s and 1.0 s from the choice onset. We then applied principal component analysis (PCA) to reduce the data to 20 principal components. We used a linear support vector machine (SVM) classifier to derive a 1-dimensional “choice axis” from the 20-dimensional data^52^. We performed 5-fold CV to determine the optimal regularization parameter (C) in the SVM. To obtain two additional axes, we applied PCA on the hyperplane orthogonal to the choice axis^53^. Similar to our body-movement decoding analysis (**Fig. 1e**), we repeated the SVM analyses 10 times in different subsampled trials and analyzed the average population trajectory of choice decoding.

### Detection of far neurons

We selected 45 sessions in which at least five neurons exhibited a cosine distance greater than 0.95 from the activity of all ANN units. We then built the HNN using these 45 sessions.

#### Quantification and statistical analysis

All analyses were performed with MATLAB (MathWorks) or Python3. We used two-sided nonparametric statistical tests. The error bars for the means are standard errors or 95% confidence intervals. The confidence intervals of sound and choice decoding were analyzed with the bootstrap method 1,000 times^54^. We used robust linear models (Python *statsmodels*) and analyzed the significance of regression coefficients with t tests (**Fig. 2e, f, S3a**). We used one-way ANOVA (Python *scypy.stats*) to compare multiple groups (**Fig. 2g, S3b**). All the statistical details are provided in the figure legends, figures, or results.

## Data and code availability

The data will be publicly available before publication. The codes are on Zenodo (https://zenodo.org/records/14749313), and will be publicly available before publication.

## Acknowledgments

This work was funded by JSPS Kakenhi (JP21H05243, JP21H00187 JP21H03492, JP22H04766, JP23K14300, JP24H02150, JP24KK0186), AMED (JP23wm0525008, JP24gm6510019, JP 24wm0625415), and the Uehara Memorial Foundation for A.F. pAAV-CKIIa-stGtACR2-FusionRed was a gift from Ofer Yizhar (Addgene plasmid #105669; http://n2t.net/addgene:105669; RRID:Addgene_105669)^55^. pAAV-CaMKIIa-mCherry was a gift from Karl Deisseroth (Addgene plasmid #114469; http://n2t.net/addgene:114469; RRID:Addgene_114469).

## Author contributions

Y.U. analyzed the data and wrote the paper. H.M. analyzed the data. S.W. collected the animal data. A.F. designed the study, analyzed the data, and wrote the paper.

## Materials & Correspondence

Should be addressed to A.F.

---

**Supplementary Figure 1.**
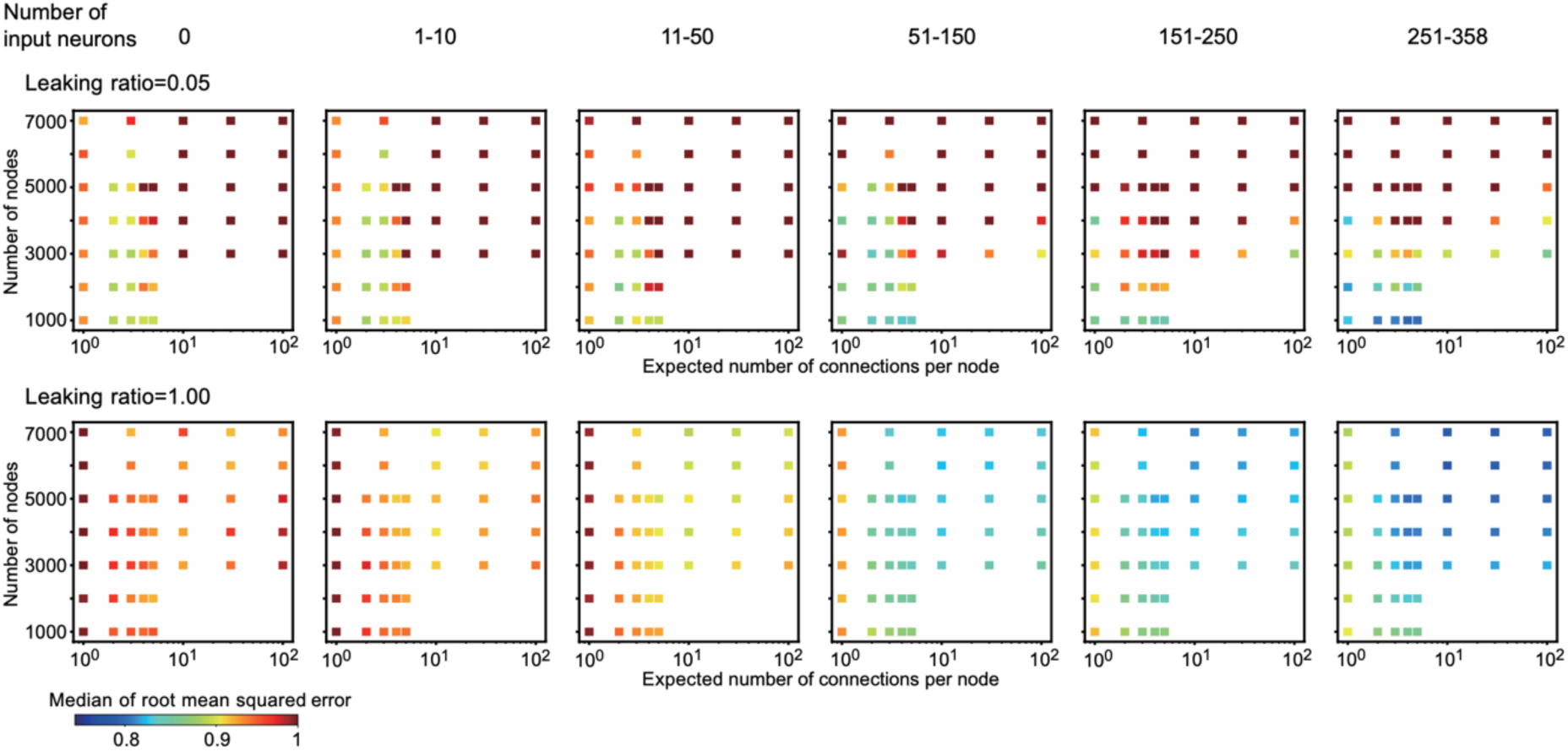
Optimal hyperparameters for reservoir computing. The performance of reservoir computing (RC) depends on the number of input neurons. The median root mean squared error (RMSE) in each parameter setting (number of nodes and expected number of connections per node) is shown with a color map. The number of sessions for each number of input neurons are as follows: 0 neurons (88 sessions), 1–10 neurons (41 sessions), 11–50 neurons (65 sessions), 51–150 neurons (39 sessions), 151–250 neurons (8 sessions), and 251–358 neurons (16 sessions). The hybrid neural network (HNN) model used data from neurons (1) in each brain region of the AC, OFC, PPC, HPC, STR, or M1 (127 sessions) and (2) in simultaneous recordings of the PPC/HPC and STR/M1 (42 sessions; 169 sessions in total). The spike data of (1) and (2) overlapped for the PPC, HPC, STR, and M1.

**Supplementary Figure 2.**
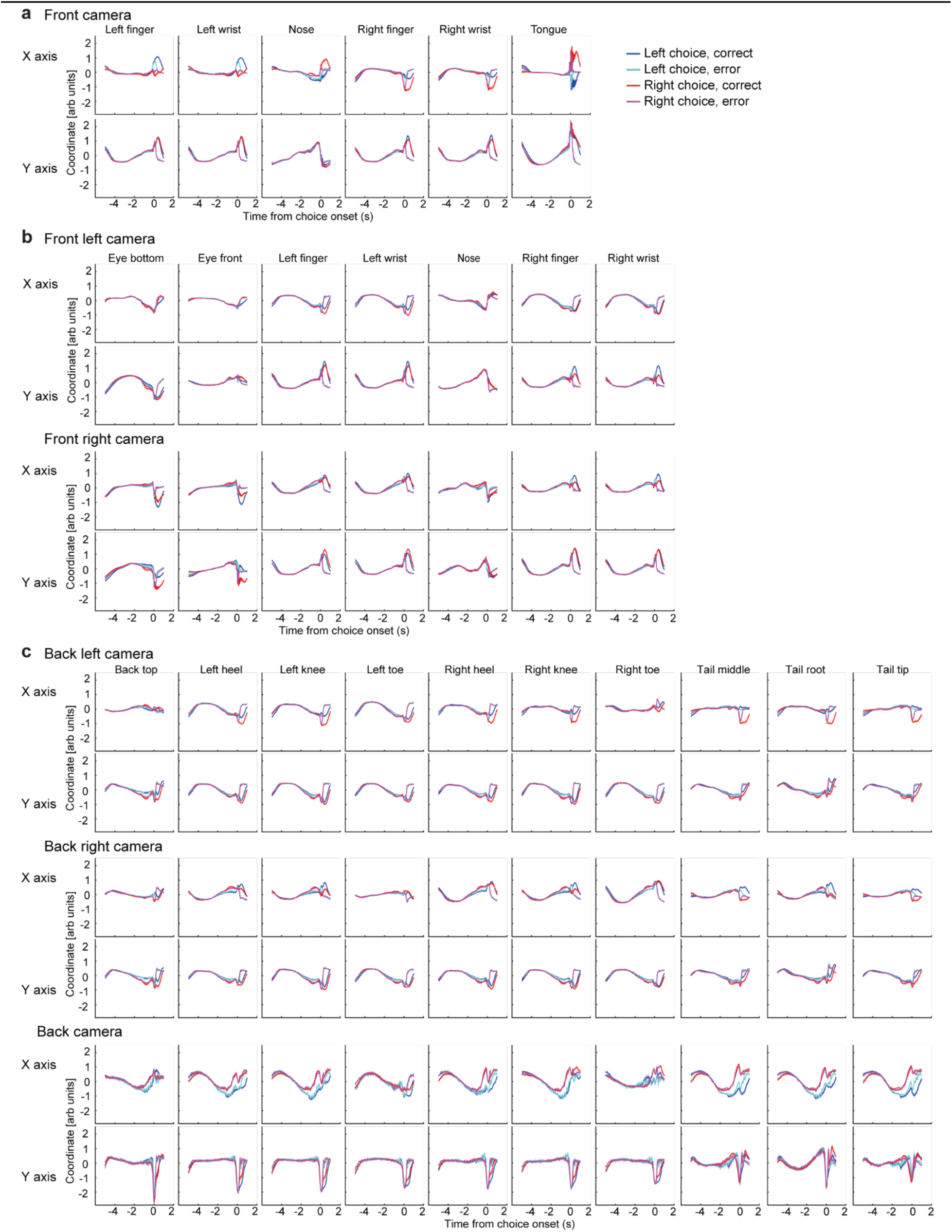
Average trajectories of body points of the mice. The average trajectories of body points from each camera are shown (**a**: front; **b**: front left and right; **c**: back left, back right, and back) (means and standard errors, 88 sessions from 6 mice).

**Supplementary Figure 3.**
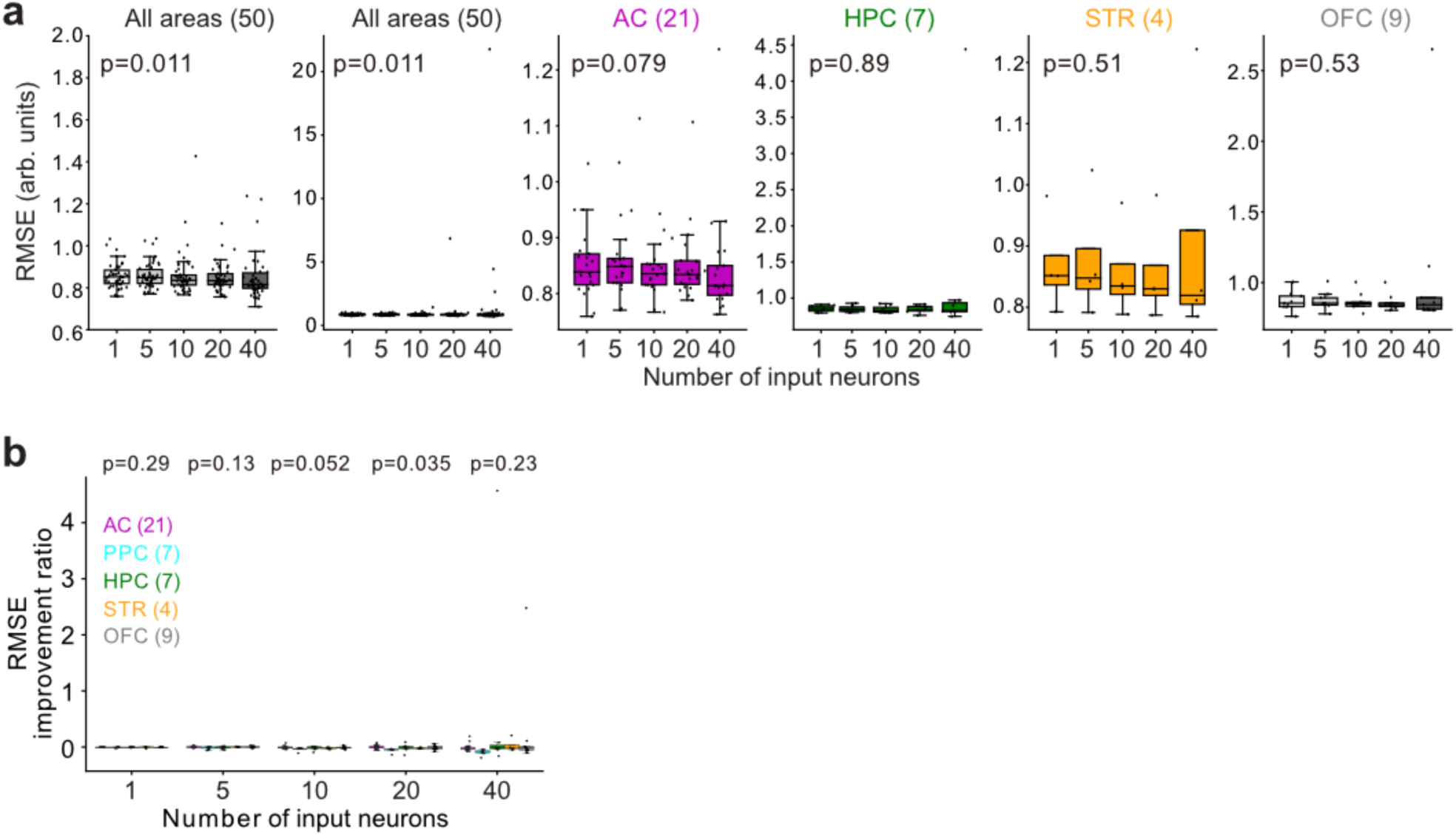
Performance of the real-cyber hybrid neural network (HNN) for all the data points. **a, b.** The data plots are the same as those in Fig. 2f and 2g, but all the data points are included. Sessions with 40 or more neurons were selected for analyses (central line and edges in the boxplots: median, 25th, and 75th percentiles; whiskers: most extreme data points inside 1.5 times the interquartile range). **a.** Relationship between the number of neurons and the performance of the HNN within sessions. All the data points in the PPC are already shown in Fig. 2f. Robust linear regression was used to analyze the significance of the slope (p values from the t test). **b.** Comparisons of RMSE improvement ratios across brain regions in each neuron number (p values from one-way ANOVA).

**Supplementary Figure 4.**
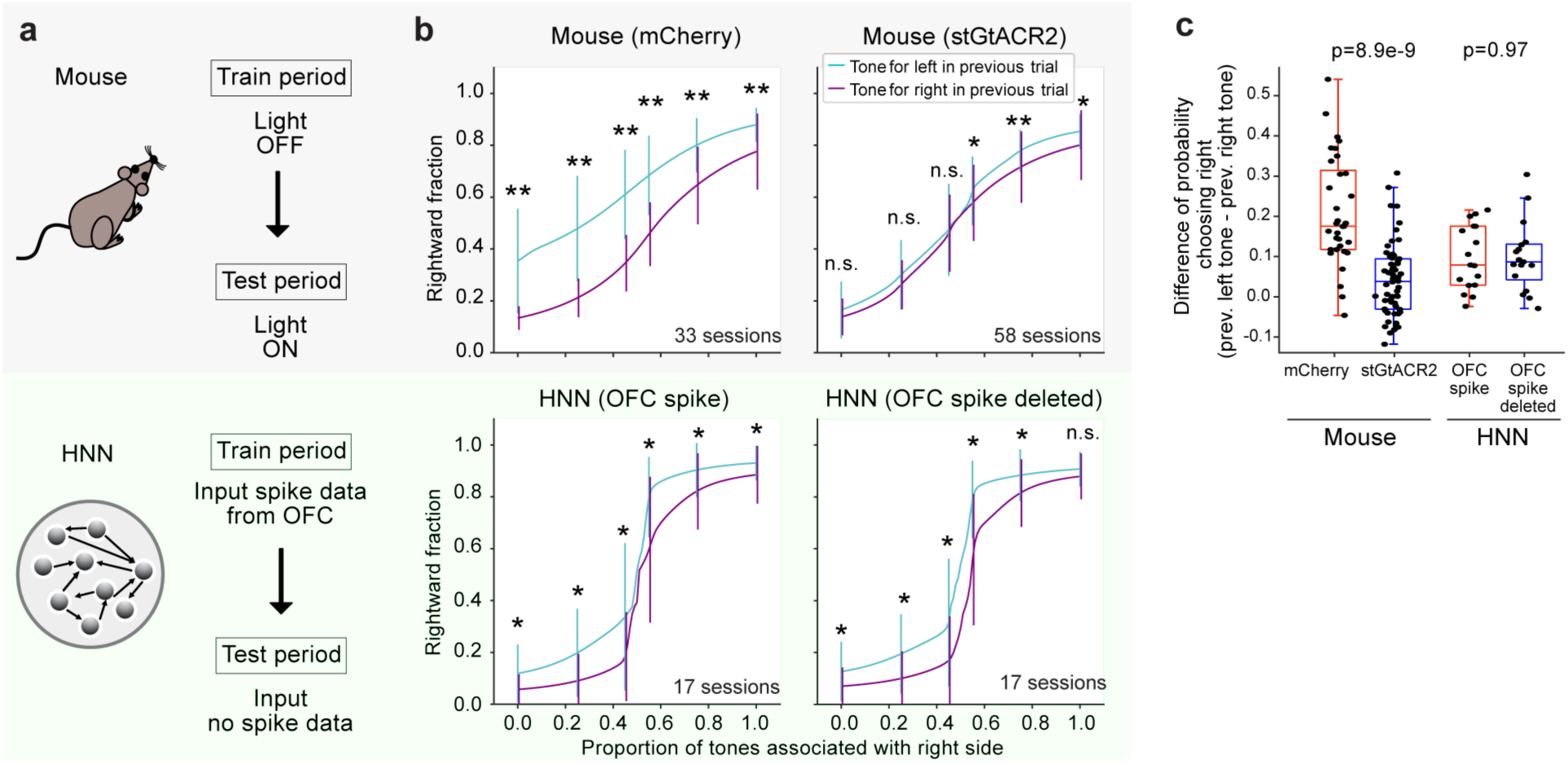
Pseudoinactivation experiment in the HNN. **a.** CBA/J mice expressing mCherry or stGtACR2 in the OFC were trained without blue-laser stimuli. The choice behaviors were tested with a blue laser (mCherry: 33 sessions from 3 mice; stGtACR2: 57 sessions from 4 mice). We trained the HNN on the activity of OFC neurons. The performance of the HNN was evaluated with and without OFC inputs (17 sessions from 4 mice). **b.** Average psychometric functions of discrete choices in mice and the HNN. For the HNN, the discrete choices were analyzed from the generated tongue movements. Model fitting of the psychometric functions was performed via maximum likelihood estimation. The probabilities of choosing right in the previous-left-tone trials and previous-right-tone trials were analyzed. Means and standard deviations of psychometric functions (Wilcoxon signed-rank test, n.s.: p ≥ 0.05, *: p < 0.05, **: p < 0.01) (mCherry: 14, 14, and 5 sessions from 3 mice; stGtACR2: 14, 14, 14, and 16 sessions from 4 mice; HNN: 2 sessions from 1 mouse and 5 sessions from 3 mice). **c.** Differences in the mean probabilities of choosing right between the previous-left and previous-right trials. We compared the differences between the mice with mCherry and stGtACR2 and between the HNN with and without OFC inputs (Mann‒Whitney U test). Central line and edges in the boxplots: median, 25th, and 75th percentiles; whiskers: most extreme data points inside 1.5 times the interquartile range.

As we reported with male CBA/J mice in our previous study^26^, we found that the choices of the control (mCherry) mice were biased according to the tone category of the previous trial. We then found that these history-dependent choice biases disappeared with the inactivation of OFC neurons with stGtACR2. The pseudoinactivation experiment in the HNN only slightly reduced the discrete choice biases.

## Supplementary Methods

### Optogenetic inhibition of the orbitofrontal cortex during the tone-frequency discrimination task

We used 10 male CBA/J mice (strain #000656; The Jackson Laboratory) that were 8-15 weeks of age at the start of behavioral training for optogenetic experiments under the alternating condition in our tone-frequency discrimination task^26^. Although the optogenetic experiments were also performed under the repeating condition in 10 mice, we did not use these data in this study because the HNN using the spike inputs during the repeating condition was only trained on 3 sessions in 1 mouse.

Similar to the animal protocol used in electrophysiological experiments, we first attached a head bar to the skull under isoflurane anesthesia (2% for induction, less than 1.5% for maintenance). We then performed a craniotomy with a diameter of 2 mm, targeting the OFC of both hemispheres (+2.6 mm AP, ± 1.4 ML, 2.1 mm depth). pAAV-CKIIa-stGtACR2-FusionRed (AAV1)^55^ or pAAV-CaMKIIa-mCherry (AAV1) (300 nL per hemisphere) was bilaterally injected into the OFC of 6 mice and 4 mice, respectively, at a speed of 60 nL/min with a Nanojector III (Drummond, Scientific Company) with a custom-made glass pipette back-filled with mineral oil. Optic fibers (Newdoon, Length 3.0 mm, NA 0.37) were bilaterally inserted into the OFC tilted 5 degrees in the medial direction. The fibers were fixed with a superbond adhesive and cyanoacrylate glue (Zap-A-Gap, PT03).

After the behavior experiments, the mice were deeply anesthetized with isoflurane (5%) and further anesthetized with a mixture of 1.5 mg/kg medetomidine, 20 mg/kg midazolam, and 25 mg/kg butorphanol. The mice were perfused with 10% formalin solution. We confirmed the fiber locations and the virus-injected areas with fixed brain slices. Due to improper locations or the absence of fiber traces, we did not use the data from 2 mice for stGtACR2 and 1 mouse for mCherry.

Similar to our previous behavioral training in mice^26^, we first trained the mice to discriminate the low- and high-category tones in the task. After the mice were able to discriminate the tones, we introduced the neutral condition, which had a transition probability of ‘p = 0.5’, to randomly present the tone categories in each trial. We then introduced the alternating condition (p = 0.9). When the percentage of correct responses for both the 100% low-tone and 100% high-tone stimuli exceeded 70% in the neutral condition, we started to deliver the blue-laser stimulus bilaterally via the optic fiber from the next session. The blue laser was 473 nm at 75 Hz (Thorlabs, CP33/M, LDC202C). Before each session, we precalibrated the laser power at the tip of the optical fibers. We set the initial laser power at 1.8 mW and gradually increased it to 4.8 mW during the neutral condition. The laser power was constant (4.8 mW) during the alternating condition. We exposed the OFC to the blue laser in all trials. Each mouse was subjected to an optogenetics experiment for at least 14 consecutive sessions, with the exception of 1 mouse with 5 sessions.

Like in the electrophysiological experiments, we analyzed the sessions in which (i) the percentage of correct responses for both the 100% low- and 100% high-tone stimuli were greater than 75% and (ii) the total reward amount in one session was at least 600 µL. For stGtACR2, we used the data from 14 sessions in 3 mice and 16 sessions in 1 mouse. For mCherry, we used the data from 14 sessions in 2 mice and 5 sessions in 1 mouse. Based on the above criteria, we excluded the data from 21 sessions in 4 mice for stGtACR2 and 7 sessions in 2 mice for mCherry.

### Comparison of optogenetic inactivation in mice and pseudoinactivation of the HNN

We used a psychometric function with a cumulative normal distribution to analyze the choice behavior of the mice during the optogenetic experiments (**Fig. S4**)^7,26^. The parameters of the psychometric function (decision threshold, stimulus sensitivity, lapse rates) were optimized to maximize the log likelihood. We also analyzed the psychometric function of the HNN. We analyzed the discrete choice in each trial according to the x-axis trajectories of the tongue movements generated by the HNN. When the tongue trajectory first exceeded one standard deviation after approaching the spout, we defined the HNN as choosing the left or right spout.

